# Tensor decomposition-Based Unsupervised Feature Extraction Applied to Single-Cell Gene Expression Analysis

**DOI:** 10.1101/684225

**Authors:** Y-h. Taguchi, Turki Turki

## Abstract

Although single cell RNA sequencing (scRNA-seq) technology is newly invented and promising one, because of lack of enough information that labels individual cells, it is hard to interpret the obtained gene expression of each cell. Because of this insufficient information available, unsupervised clustering, e.g., t-Distributed Stochastic Neighbor Embedding and Uniform Manifold Approximation and Projection, is usually employed to obtain low dimensional embedding that can help to understand cell-cell relationship. One possible drawback of this strategy is that the outcome is highly dependent upon genes selected for the usage of clustering. In order to fulfill this requirement, there are many methods that performed unsupervised gene selection. In this study, a tensor decomposition (TD) based unsupervised feature extraction (FE) was applied to the integration of two scRNA-seq expression profiles that measure human and mouse midbrain development. TD based unsupervised FE could not only select coincident genes between human and mouse, but also biologically reliable genes. Coincidence between two species as well as biological reliability of selected genes is increased compared with principal component analysis (PCA) based FE applied to the same data set in the previous study. Since PCA based unsupervised FE outperformed other three popular unsupervised gene selection methods, highly variable genes, bimodal genes and dpFeature, TD based unsupervised FE can do so as well. In addition to this, ten transcription factors (TFs) that might regulate selected genes and might contribute to midbrain development are identified. These ten TFs, BHLHE40, EGR1, GABPA, IRF3, PPARG, REST, RFX5, STAT3, TCF7L2, and ZBTB33, were previously reported to be related to brain functions and diseases. TD based unsupervised FE is a promising method to integrate two scRNA-seq profiles effectively.

## 1 INTRODUCTION

Single cell RNA sequencing (scRNA-seq) (17) is a newly invented technology that enables us to measure amount of RNA in single cell basis. In spite of its promising potential, it is not easy to interpret the measurements. The primary reason of this difficulty is the lack of sufficient information that characterizes individual cells. In contrast to the huge number of cells measured, which is often as many as several thousands, the number of labeling is limited, e.g., measurement of conditions as well as the amount of expression of key genes measured by fluorescence-activated cell sorting, whose number is typically as little as tens. This prevents us from selecting genes that characterize the individual cell properties.

In order to deal with samples without suitable numbers of labelling, unsupervised method is frequently used, since it does not make use of labeling information directly. K-means clustering as well as hierarchical clustering are the popular methodology that are often applied to gene expression analysis. The popular clustering methods specifically applied to scRNA-seq is tSNE (52) or UMAP (9), which is known to be useful to get low dimensional embedding of a set of cells. In spite of that, the obtained clusters are highly dependent upon genes used for clustering. Thus, the next issue is, without labeling (i.e., pre-knowledge), to select genes that might be biologically meaningful.

The various unsupervised gene selection methods applicable to scRNA-seq were invented, e.g., highly variable genes, bimodal genes and dpFeature and principal component analysis (PCA) based unsupervised feature extraction (FE) (50, 34, 49, 42, 25, 31, 27, 40, 47, 4, 5, 44, 45, 13, 12, 11, 51, 39, 43, 22, 23, 24, 26, 41, 48). Chen et al. (2) recently compared genes selected by these methods and concluded that the genes selected are very diverse and have their own (unique) biological features. In this sense, it is required to invent more advanced unsupervised gene selection methods that can select more biologically relevant genes.

In this paper, we propose the application of tensor decomposition (TD) based unsupervised FE (37, 46, 35, 36, 30, 32, 29, 33, 28). It is an advanced method of PCA based unsupervised FE, to scRNA-seq analysis. For more details about PCA based unsupervised FE and TD based unsupervised FE, see the recently published book (38). Especially focusing on the integration of two scRNA-seq profiles, the advantages of TD based unsupervised FE when compared with PCA based unsupervised FE is as follows; The former can integrate more than two gene expressions prior to the analysis while the latter can only integrate the results obtained by applying the method to individual data sets.

In the following, based on the previous study (34) where PCA based unsupervised FE was employed, we try to integrate human and mouse midbrain development gene expression profiles to obtain key genes that contribute to this process, by applying TD based unsupervised FE. It turned out that TD based unsupervised FE can identify biologically more relevant and more common genes between human and mouse than PCA based unsupervised FE that outperformed other methods compared with.

## 2 METHODS AND MATERIALS

### 2.1 scRNA-seq data

#### 2.1.1 Midbrain development of humans and mice

The first scRNA-seq data used in this study was downloaded from gene expression omnibus (GEO) under the GEO ID GSE76381; the files named “GSE76381 EmbryoMoleculeCounts.cef.txt.gz” (for human) and “SE76381 MouseEmbryoMoleculeCounts.cef.txt.gz” (for mouse) were downloaded. These two gene expression profiles generated from scRNA-seq data set; One represents human embryo ventral midbrain cells between 6 and 11 weeks of gestation (287 cells for six weeks, 131 cells for seven weeks, 331 cells for eight weeks, 322 cells for nine weeks, 509 cells for ten weeks, and 397 cells for eleven weeks, in total 1977 cells). Another is a set of mouse ventral midbrain cells at six developmental stages between E11.5 to E18.5 (349 cells for E11.5, 350 cells for E12.5, 345 cells for E13.5, 308 cells for E14.5, 356 cells for E15.5, 142 cells for E18.5 and 57 cells for unknown, in total 1907 cells).

#### 2.1.2 Mouse hypothalamus with and without acute formalin stress

The second scRNA-seq data used in this study were downloaded from GEO under GEOID GSE74672; the file named “GSE74672 expressed mols with classes.xlsx.gz” was downloaded. It is generated from snRNA-seq data set that measures mouse hypothalamus with and without acute formalin stress. Various mete-data, which is included in the first eleven rows of data set, are available. The meta-data available includes, sex, age, cell types (astrocytes, endothelial, ependymal, microglia, neurons, oligos and vsm) and control vs stressed samples and so on.

### 2.2 TD based unsupervised FE

#### 2.2.1 Midbrain development of humans and mice

TD based unsupervised FE is recently proposed methods successfully applied to various biological problems. TD based unsupervised FE can be used for integration of multiple measurements applied to the common set of genes. Suppose *x*_*ij*_ ∈ ℝ^*N*×*M*^ and *x*_*ik*_ ∈ ℝ^*N*×*K*^ are the *i*th expression of *j*th and *k*th cells under the two distinct conditions (in the present study, they are human and mouse), respectively. Then 3-mode tensor, *x*_*ijk*_ ∈ ℝ^*N*×*M*×*K*^, where *N* (= 13889) is total number of common genes between human and mouse, which share gene symbols, between human and mouse, *M* (= 1977) is the number of human cells and *K*(= 1907) is total number of mouse cells, is defined as

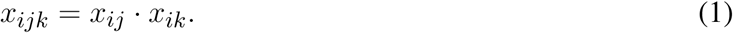

It is Case I Type II tensor (32). Here Case I means that tensor is generated such that two matrices share the samples while Type II means that summation is taken over as in eq. (2). On the other hand, tensor before taking summation as in eq. (1) is Tupe I. Since it is too large to be decomposed, we further transform it into type II tensor as

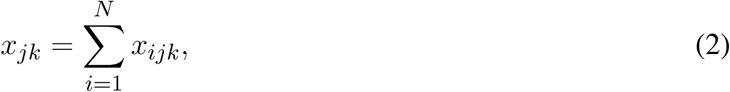

where *x*_*jk*_ ∈ ℝ^*M*×*K*^ is now not a tensor but a matrix. In this case, TD is equivalent to singular value decomposition (SVD). After applying SVD to *x*_*jk*_, we get SVD,

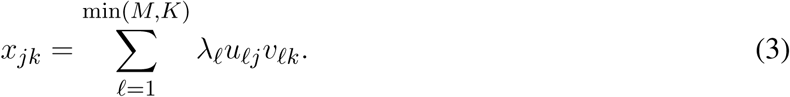

where *u*_*ℓj*_ ∈ ℝ^*M*×*M*^ and *v*_*ℓk*_ ∈ ℝ^*K*×*K*^ are singular value vectors attributed to cells of human scRNA-seq and those of mouse scRNA-seq, respectively.

Singular value vectors attributed to genes of human and mouse scRNA-seq, *u*_*ℓi*_ ∈ ℝ^*N*×*M*^ and *v*_*ℓi*_ ∈ ℝ^*N*×*K*^, are defined as

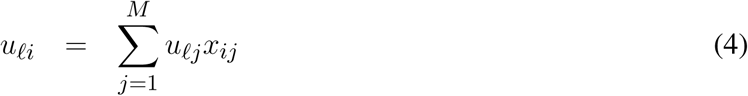

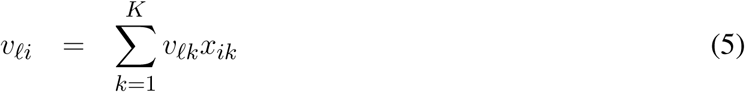

respectively.

In order to find genes associated with biological functions, we need to select *u*_*ℓj*_ and *v*_*ℓk*_ which are coincident with biological meaning. In this study, we employ time points of measurements as biological meanings. In other words, we seek genes associated with time development. Since we would like to find any kind of time dependence, we simply deal with time points as un-ordered labelling. Thus, we apply categorical regression

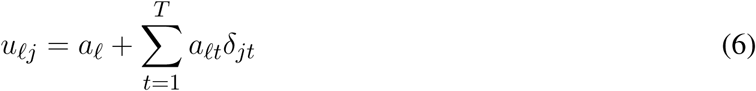

(T=6; t=1 to T correspond to 6, 7, 8, 9, 10 and 11 weeks, see Methods and Materials) or

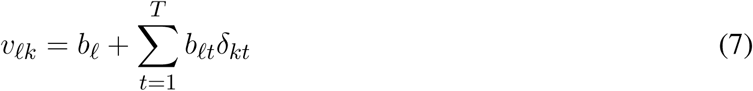

(T=7; t=1 to T correspond to E11.5, E12.5, E13.5, E14.5, F15.5, E18.5 and unknown, see Methods and Materials) where *δ*_*jt*_(*δ*_*kt*_) = 1 when the *j*th(*k*th) cell is taken from the *t*th time point otherwise *δ*_*jt*_(*δ*_*kt*_) = 0. *a*_*ℓ*_, *a*_*ℓt*_, *b*_*ℓ*_ and *b*_*ℓt*_ are the regression coefficients.

*P* -values are attributed to *ℓ*th singular value vectors using the above categorical regression (lm function in R (15) is used to compute *P* -values). *P* -values attributed to singular value vectors are corrected by BH criterion (1). Singular value vectors associated with corrected *P* -values less than 0.01 are selected for the download analysis. Hereafter, the set of selected singular value vectors of human and mouse are denoted as 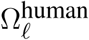 and 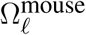, respectively.

*P* -values are attributed to genes with assuming *χ*^2^ distribution for the gene singular value vectors, *u*_*ℓi*_ and *v*_*ℓi*_, corresponding to the cell singular value vectors selected by categorical regression as

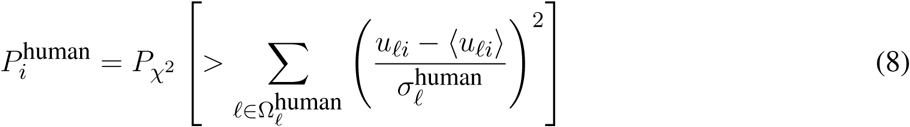

for human genes and

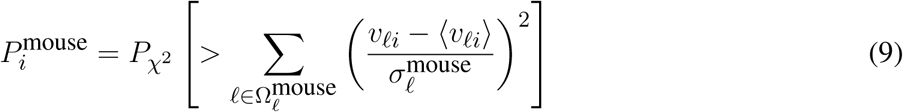

for mouse genes, respectively. Here

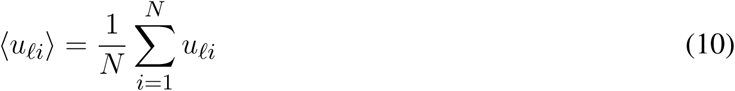

and

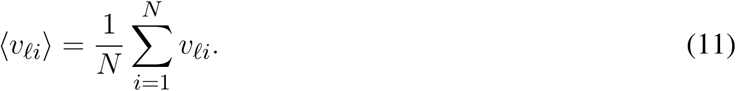

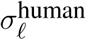 and 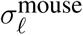 are the standard deviations of *ℓ*th gene singular value vectors for human and mouse respectively, 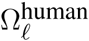 and 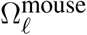 are sets of *ℓ*s selected by categorical regression for human (eq. (6)) and mouse (eq.(7)), respectively, and 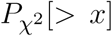 is the cumulative probability of *χ*^2^ distribution when the argument takes values larger than *x*. 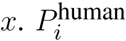 and 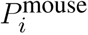 are corrected by BH criterion and genes associated with corrected *P* -values less than 0.01 are selected.

#### 2.2.2 Mouse hypothalamus with and without acute formalin stress

The application of TD based unsupervised FE to mouse hypothalamus is quit similar to that to mouse and human midbrain. There are also two matrices, *x*_*ij*_ ∈ ℝ^*N*×*M*^ and *x*_*ik*_ ∈ ℝ^*N*×*K*^ that correspond to *i*th expression of *j*th and *k*th cells under the two distinct conditions (in the present case, they are without and with acute formalin stress, respectively); *N* = 24341, *M* = 1785 and *K* = 1096. Type I Case II tensor, *x*_*jk*_, was also generated using eqs. (1) and (2) and SVD was applied to *x*_*jk*_ as eq. (3). Then singular vale vectors attributed to genes of samples without and with acute formalin stress, *u*_*ℓi*_ and *v*_*ℓi*_ were computed by eqs. (4) and (5). We alos applied categorical regressions to *u*_*ℓi*_ and *v*_*ℓi*_, although categories considered here are not time points but cell types. Then categorical regressions applied to *u*_*ℓi*_ and *v*_*ℓi*_ in mouse hypothalamus without and with acute formalin stress are

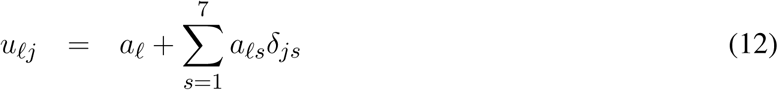

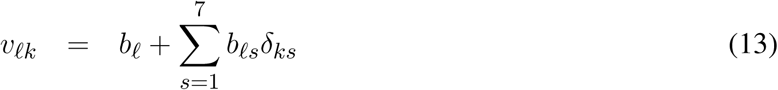

where *s* stands for one of seven cell types mentioned in §2.1.2 *and δ*_*js*_(*δ*_*ks*_) = 1 when the *j*th(*k*th) cell is taken from the *s*th cell types otherwise *δ*_*jt*_(*δ*_*kt*_) = 0. Table 1 lists the number of cells in these categories. The remaining procedures to select genes associated with identified cell type dependency is exactly the same as those in midbrain case.

**Table 1.** The number of cells that belong to either without and with acute formalin stress or cell types

| cell types | without | with |
| --- | --- | --- |
|  | acute stress |  |
| astrocytes | 135 | 132 |
| endothelial | 169 | 71 |
| ependymal | 211 | 145 |
| microglia | 34 | 14 |
| neurons | 628 | 270 |
| oligos | 570 | 431 |
| vsm | 38 | 33 |

### 2.3 Enrichment analyses

Various enrichment analysis methods are performed with separately uploading selected human and mouse gene symbols, or genes selected commonly between samples without and samples with acute formalin stress, to Enrichr (6).

## 3 RESULTS

### 3.1 Midbrain development of humans and mice

As a result, following the procedure described in Methods and Materials, we identified 55 and 44 singular value vectors attributed to cells, *u*_*ℓj*_s and *v*_*ℓk*_s, for human and mouse, respectively. One possible validation of selected *u*_*ℓj*_s and *v*_*ℓk*_s are coincidence. Although cells measured are not related between human and mouse at all, if SVD works well, corresponding singular value vectors (i.e., *u*_*ℓj*_ and *v*_*ℓk*_ sharing the same *ℓ*s) attributed to cells should share something biological, e.g., time dependence. This suggests that it is more likely that corresponding singular value vectors attributed to cells, *u*_*ℓj*_ and *v*_*ℓk*_, are simultaneously associated with significant *P* -vales computed by categorical regression. As expected, they are highly significantly correlated. Table 2 shows confusion matrix of the coincidence of selected singular value vectors between human and mouse. For human cells, only top 1907 singular value vectors among all 1977 singular value vectors are considered, since total number of singular value vectors attributed to mouse cells is 1907. selected: corrected *P* -values, computed with regression analysis eqs. (6) and (7), are less than 0.01, not selected: otherwise. Odds ratio is as many as 227 and *P* -values computed by Fisher’s exact test is 1.44× 10^−44^.

**Table 2.** Confusion matrix of coincidence between selected 55 singular value vectors selected among all 1977 singular value vectors, *u*_*ℓj*_, attributed to human cells and 44 singular value vectors selected among all 1907 singular value vectors, *v*_*ℓk*_, attributed to mouse cells.

|  |  | human |  |
| --- | --- | --- | --- |
|  |  | not selected | selected |
| mouse | not selected | 1833 | 12 |
|  | selected | 23 | 32 |

Figure 1 shows the coincidence of selected singular value vectors between human and mouse. Singular value vectors with smaller *ℓ*s, i.e., with more contributions, are more likely selected and coincident between human and mouse. This can be the side evidence that guarantees that TD based unsupervised FE successfully integrated human and mouse scRNA-seq data.

**Figure 1.**
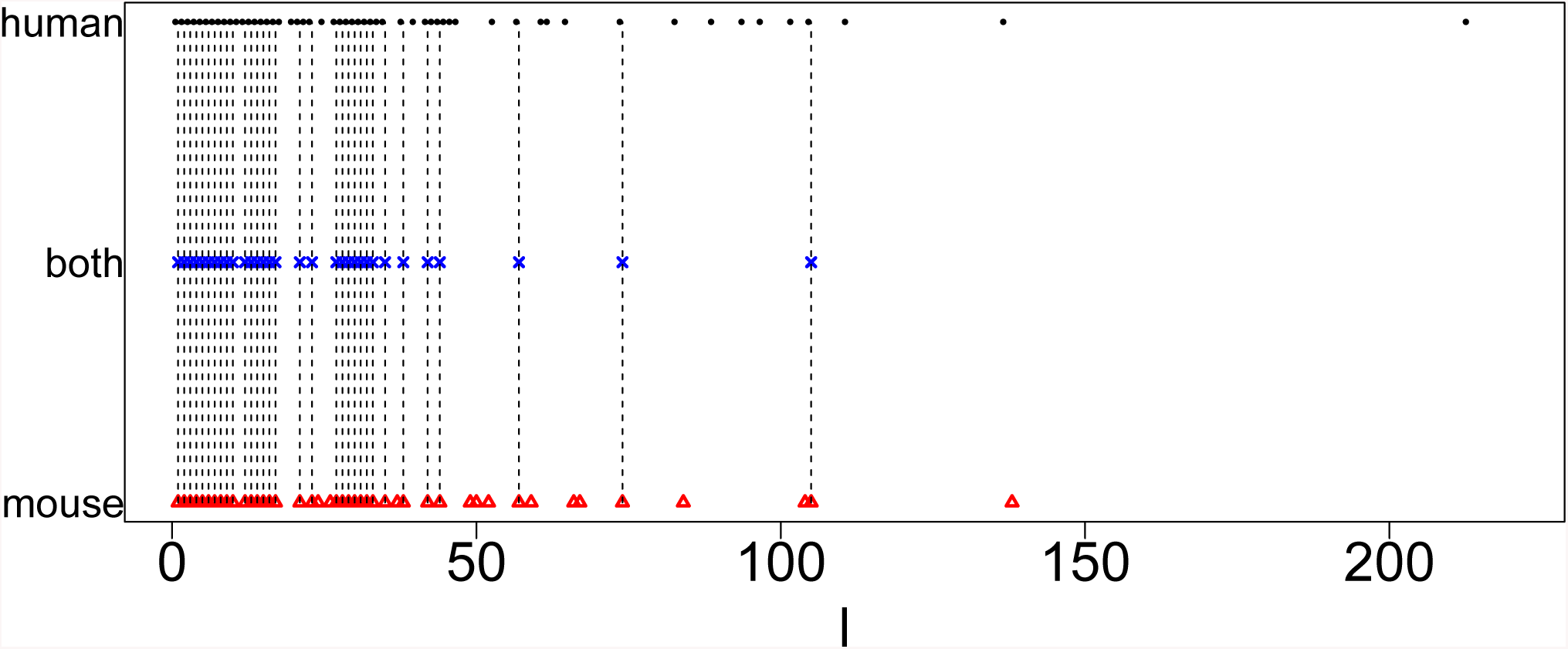
Coincidence between singular value vectors shown in Table 2. Horizontal axis: singular value vector numbering *ℓ*. Black open circles: *ℓ*s selected for human, blue crosses: *ℓ*s selected for both human and mouse, red open triangles: *ℓ*s selected for mouse. Vertical black broken lines connect *ℓ*s selected for both human and mouse.

Next, we selected genes with following the procedures described in Methods and Materials. The first validation of selected genes is the coincidence between human and mouse. In my previous study (34), more number of common genes were selected by PCA based unsupervised FE than other methods compared, i.e., highly variables genes, bimodal genes and dpFeature. Table 3 shows the confusion matrix that describes the coincidence of selected genes between human and mouse. Odds ratio is as large as 133 and *P* -value is 0 (i.e., less than numerical accuracy), which is significantly better than coincidence of selected genes between human and mouse (53 common genes between 116 genes selected for human and 118 genes selected mouse), previously achieved by PCA based unsupervised FE (34), which outperformed other methods, i.e., highly variable genes, bimodal genes and dpFeature.

**Table 3.** Confusion matrix of coincidence between selected 456 genes for human and selected 505 genes for mouse among all 13384 common genes. Selected: corrected *P* -values, computed with *χ*^2^ distribution eqs. (8) and (9), are less than 0.01, not selected: otherwise. Odds ratio is as many as 133 and *P* -values computed by Fisher’s exact test is 0 (i.e. less than numerical accuracy).

|  |  | human |  |
| --- | --- | --- | --- |
|  |  | not selected | selected |
| mouse | not selected | 13233 | 151 |
|  | selected | 200 | 305 |

On the other hands, most of genes selected by PCA based unsupervised FE in the previous study (34) are included in the genes selected by TD based unsupervised FE in the present study. One hundred and two genes are selected by TD based unsupervised FE among 116 human genes selected by PCA based unsupervised FE in the previous study (34) while 91 genes are selected by TD based unsupervised FE among 118 mouse genes by PCA based unsupervised FE. Thus, TD based unsupervised FE is quite consistent with PCA based unsupervised FE.

Biological significance tested by enrichment analysis is further enhanced. Most remarkable advances achieved by TD based unsupervised FE is “Allen Brain Atlas”, to which only downregulated genes were enriched in the previous study (34). As can be seen in Table 4, now many enrichment associated with upregulated genes. In addition to this, most of top ranked five terms are related to paraventricular, which is adjusted to midbrain. This suggests that TD based unsupervised FE successfully identified genes related to midbrain.

**Table 4.**
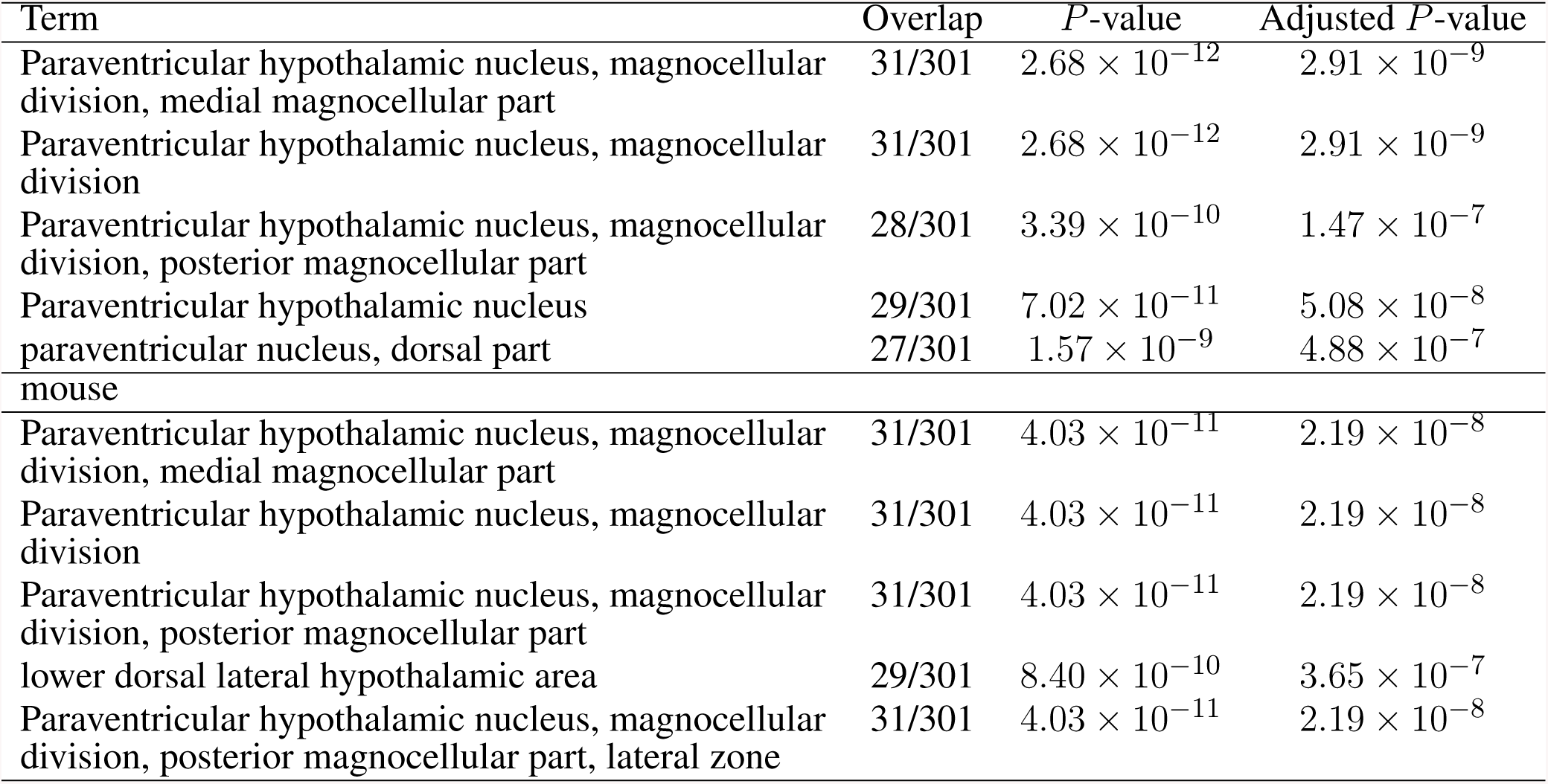
Five top ranked terms from “Allen Brain Atlas up” by Enrichr for selected 456 human genes and 505 mouse genes.

| human |  |  |  |  |
| --- | --- | --- | --- | --- |
| Term | Overlap | $P$-value | Adjusted $P$-value | |
| Paraventricular hypothalamic nucleus, magnocellular division, medial magnocellular part | 31/301 | $2.68 \times 10^{-12}$ | $2.91 \times 10^{-9}$ | |
| Paraventricular hypothalamic nucleus, magnocellular division | 31/301 | $2.68 \times 10^{-12}$ | $2.91 \times 10^{-9}$ | |
| Paraventricular hypothalamic nucleus, magnocellular division, posterior magnocellular part | 28/301 | $3.39 \times 10^{-10}$ | $1.47 \times 10^{-7}$ | |
| Paraventricular hypothalamic nucleus | 29/301 | $7.02 \times 10^{-11}$ | $5.08 \times 10^{-8}$ | |
| paraventricular nucleus, dorsal part | 27/301 | $1.57 \times 10^{-9}$ | $4.88 \times 10^{-7}$ | |
| mouse |  |  |  |  |
| Paraventricular hypothalamic nucleus, magnocellular division, medial magnocellular part | 31/301 | $4.03 \times 10^{-11}$ | $2.19 \times 10^{-8}$ | |
| Paraventricular hypothalamic nucleus, magnocellular division | 31/301 | $4.03 \times 10^{-11}$ | $2.19 \times 10^{-8}$ | |
| Paraventricular hypothalamic nucleus, magnocellular division, posterior magnocellular part | 31/301 | $4.03 \times 10^{-11}$ | $2.19 \times 10^{-8}$ | |
| lower dorsal lateral hypothalamic area | 29/301 | $8.40 \times 10^{-10}$ | $3.65 \times 10^{-7}$ | |
| Paraventricular hypothalamic nucleus, magnocellular division, posterior magnocellular part, lateral zone | 31/301 | $4.03 \times 10^{-11}$ | $2.19 \times 10^{-8}$ | |

In addition to this, “Jensen TISSUES” (Table 5) for Embryonic brain is highly enhanced (i.e., more significant (smaller) *P* -values ∼ 10^−100^, which were as large as 10^−10^ to 10^−20^ in the previous study (34)). On the other hand, “ARCHS4 tissues” also strongly supports the biological reliability of selected genes (Table 6). The term “MIDBRAIN” is enriched highly and it is top ranked for both human and mouse.

**Table 5.**
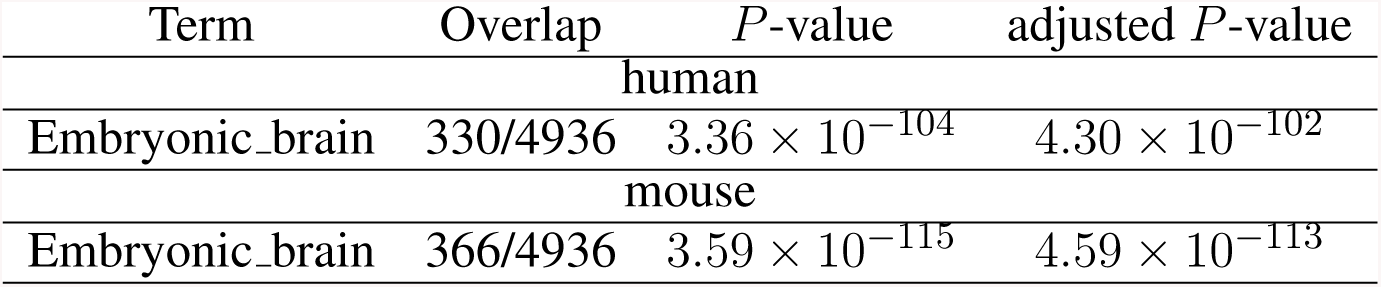
Enrichment of Embryonic brain by “JENSEN TISSUES” in Enrichr

| Term | Overlap | *P*-value | adjusted *P*-value |
| --- | --- | --- | --- |
| human |  |  |  |
| Embryonic_brain | 330/4936 | $3.36 \times 10^{-104}$ | $4.30 \times 10^{-102}$ |
| mouse |  |  |  |
| Embryonic_brain | 366/4936 | $3.59 \times 10^{-115}$ | $4.59 \times 10^{-113}$ |

**Table 6.** Enrichment of Embryonic brain by “ARCHS4 Tissues” in Enrichr

| Term | Overlap | *P*-value | adjusted *P*-value |
| --- | --- | --- | --- |
| human |  |  |  |
| MIDBRAIN | 248/2316 | $1.02 \times 10^{-129}$ | $1.11 \times 10^{-127}$ |
| mouse |  |  |  |
| MIDBRAIN | 248/2316 | $1.44 \times 10^{-99}$ | $1.56 \times 10^{-97}$ |

There are some brain related enrichment found in other categories, although they are not strong enough compared with the above three. Brain related terms in “GTEx Tissue Sample Gene Expression Profiles up” (Table 7) are also enhanced for mouse brain (top three terms are brain), although no brain terms are enriched within top five raked terms for human (This discrepancy cannot be understood at the moment). Contrary, brain related terms in “MGI Mammalian Phenotype 2017” (Table 8) are enhanced for human brain (fourth and fifth ranked), although no brain terms are enriched within top five raked terms for mouse (This discrepancy also cannot be understood at the moment). The above observations suggest that TD based unsupervised FE could identify genes related to mouse and human embryonic midbrain.

**Table 7.**
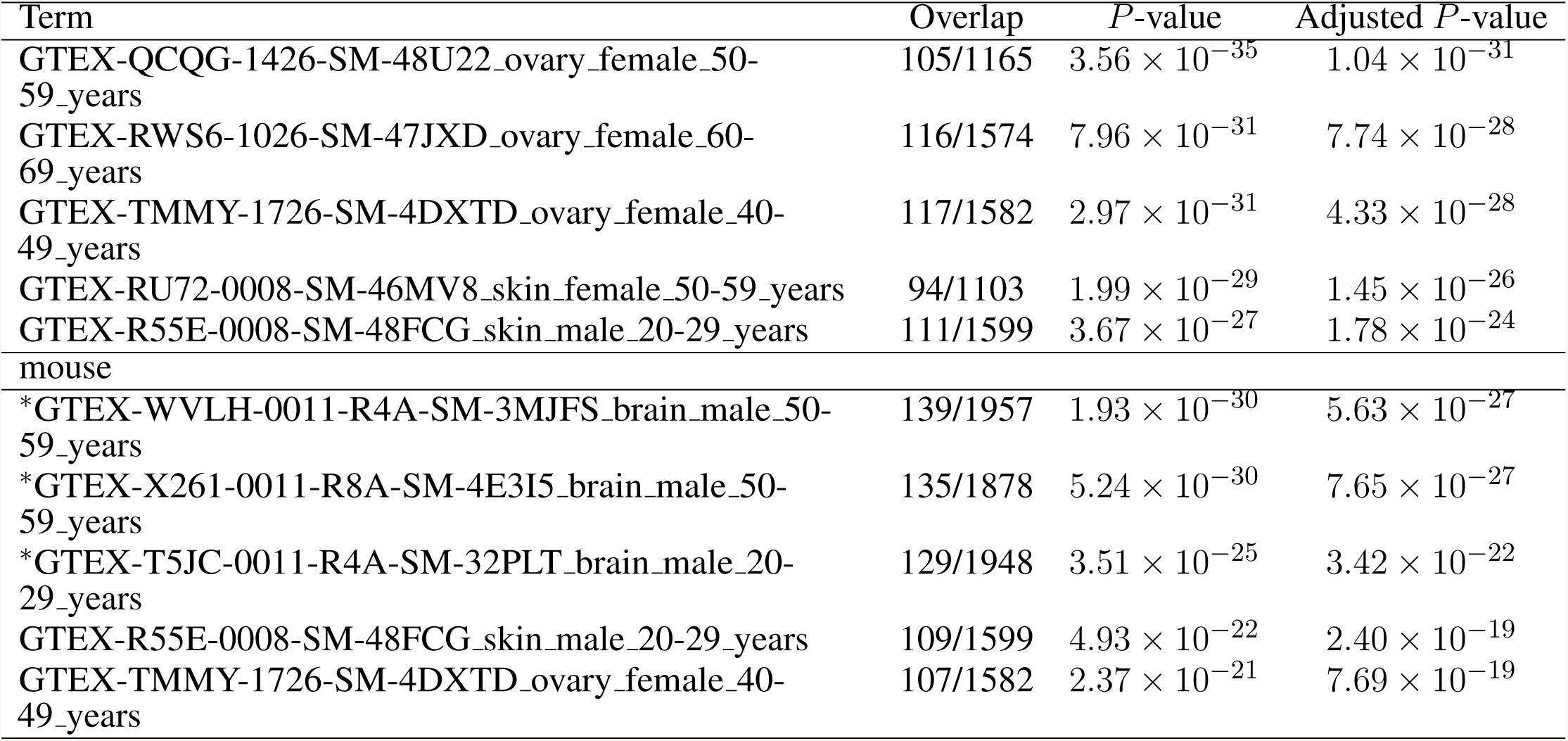
Five top ranked terms from “GTEx Tissue Sample Gene Expression Profiles up” by Enrichr for selected 456 human genes and 505 mouse genes. Brain related terms are asterisked.

**Table 8.**
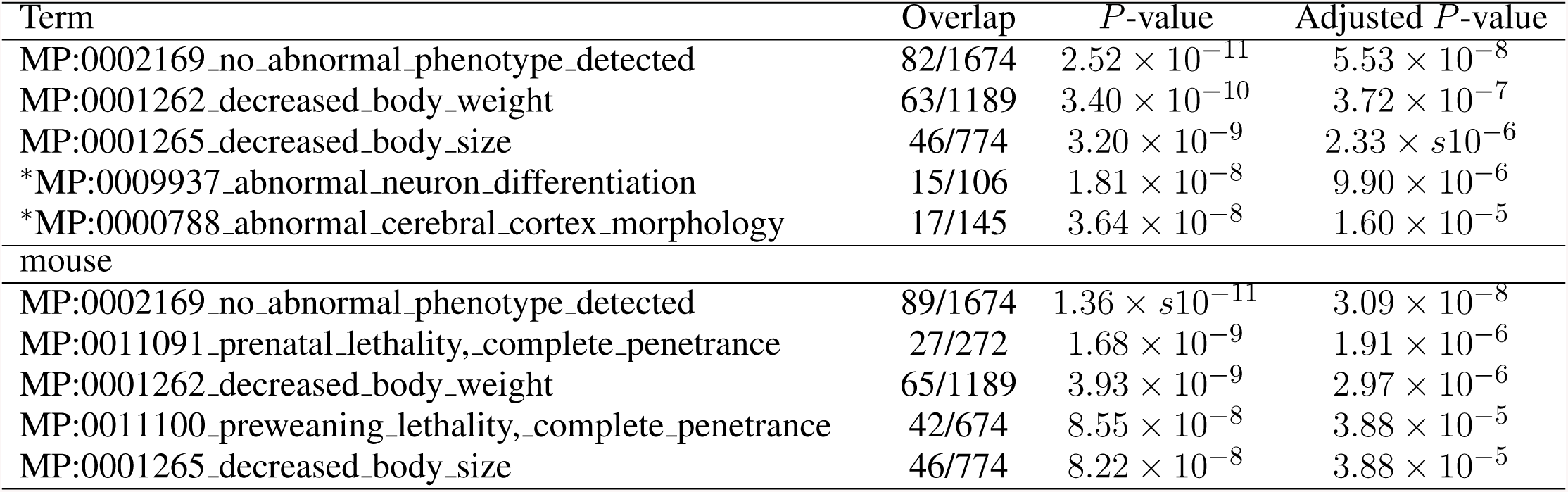
Five top ranked terms from “MGI Mammalian Phenotype 2017” by Enrichr for selected 456 human genes and 505 mouse genes. Brain related terms are asterisked.

We also uploaded selected 456 human genes and 505 mouse genes to STRING server (21) that evaluates protein-protein interaction (PPI) enrichment. 7488 PPI are reported among 456 human genes while expected number of PPI is as small as 3524 (*P* -value is less than 1× 10^−6^). 6788 PPI are reported among 505 mouse genes while expected number of PPI is as small as 3290 (*P* -value is again less than 1× 10^−6^). Thus, TD based unsupervised FE can successfully identify significantly interacting protein coding genes.

Finally, we checked if transcription factor (TF) that targets selected genes are common between human and mouse (Table 9). These TFs are associated with adjusted *P* -values less than 0.01 in “ENCODE and ChEA Consensus TFs from ChIP-X” of Enrichr. They are highly overlapped between human and mouse (There are ten common TFs between 16 TFs found in human and 24 TFs found in mouse). Although selected TFs are very distinct from those in the previous study (34), they are highly interrelated with each other (see below). These TFs are up-loaded to the regnetworkweb server (8) and TF networks shown in Figure 2 are identified. Clearly, even in partial, these TFs interact highly with each other.

**Table 9.** TFs enriched in “ENCODE and ChEA Consensus TFs from ChIP-X” by Enrichr for human and mouse. Bold TFs are common.

|  |  |
| --- | --- |
| human | BCL3, **BHLHE40**, **EGR1**, **GABPA**, **IRF3**, **PPARG**, **REST**, **RFX5**, SP1, SP2, SRF, **STAT3**, **TCF7L2**, TRIM28, TRIM28, **ZBTB33**, |
| mouse | **BHLHE40**, CTCF, E2F4, E2F6, **EGR1**, ESR1, ETS1, FLI1, **GABPA**, **IRF3**, NFIC, NRF1, **PPARG**, RCOR1, **REST**, **RFX5**, SPI1, **STAT3**, **TCF7L2**, USF1, USF2, YY1, **ZBTB33**, ZNF384, |

**Figure 2.**
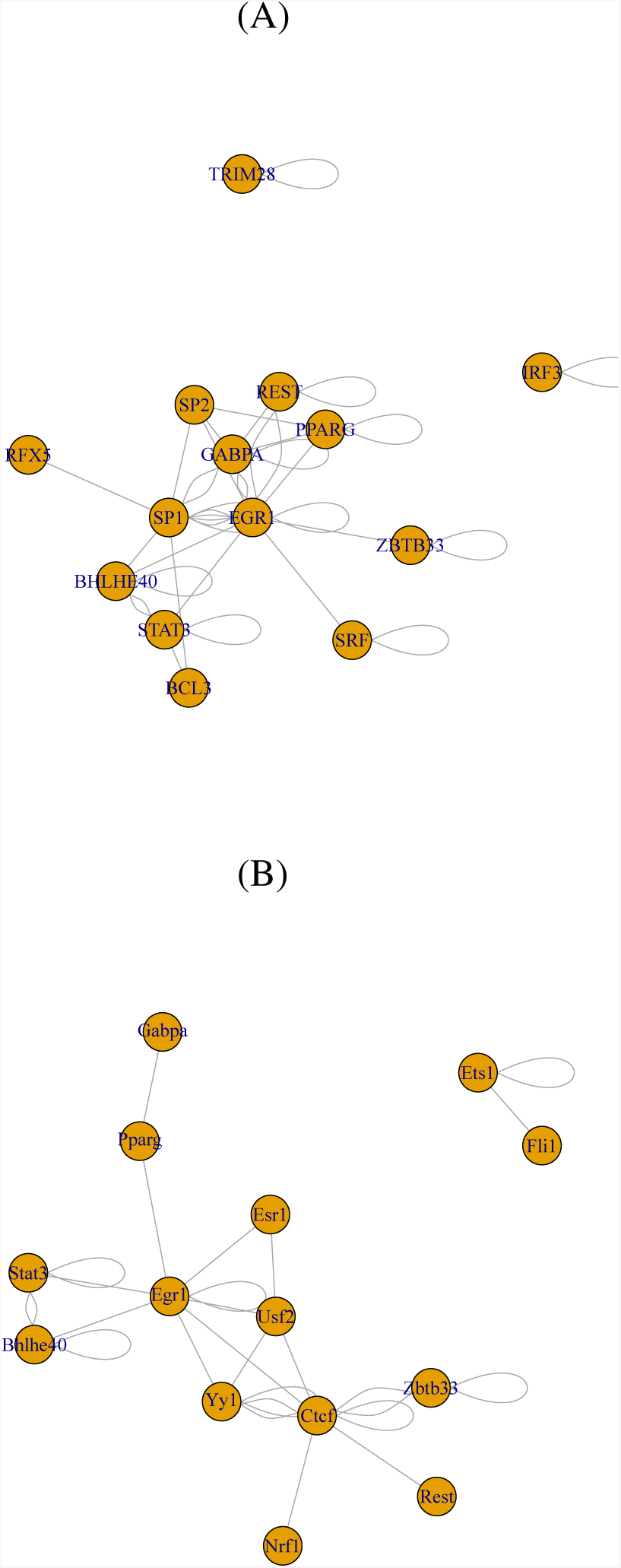
TF network identified by regnetworkweb for TFs in Table 9. (A) human, (B) mouse.

We also checked if commonly selected ten TFs (bold in Table 9) are related to brains. Lack of BHLHE40 was found to result in brain malfunction (3). The function of EGR1 was found in embryonic rat brain (54). GABPA is essential for human cognition (16). IRF3 is related to brain disease (18). PPAR, which PPARG belongs to, is believed to be therapeutic target of neurodegenerative diseases (53). REST is a master regulator of neurogenesis (10). RFX5 is known to be expressive in fetal brain (20). STAT3 promotes brain metastasis (14). TCF7L2 regulated brain gene expression (19). ZBTB33 affects the mouse behaviour through regulating brain gene expression (7). Thus all commonly selected ten TFs are related to brains.

### 3.2 Mouse hypothalamus with and without acute formalin stress

Although the effectiveness of the proposed strategy toward scRNA-seq is obvious in the results shown in the previous subsection, one might wonder if it is accidental. In order to dispel such doubts, we apply TD based unsupervised FE to yet another scRNA-seq data set: mouse hypothalamus with and without acute formalin stress. Contrary to the data set analyzed in the previous subsection where very distant two data sets were anaylised, the data sets analyzed here are very close to each other. Both data sets are taken from the same tissue of mouse, hypothalamus. The only difference is if they are stressed by formalin dope or not. The motivation why we here specifically apply TD based unsupervised FE to two close data sets is as follows; when two data sets are too close, it might be difficult to identify which genes are commonly altered by additional condition, in this case, the dependence upon cell types, because all genes might behave equally between two. Thus, it is not a bad idead to check if TD based unsupervised FE can work well not only when very distant data sets are analyzed but also very close data set is analyzed.

With following the procedure described in the materials and methods, we identified 30 and 24 singular value vectors attributed to cells, *u*_*ℓj*_s and *v*_*ℓk*_s, without and with acute formalin stress, respectively. We again applied Fisher’s exact test (Table 10). Although odds ratio is ten times larger than that in Table 2, *P* -value is even smaller than that in Table 2; This suggests that TD based unsupervised FE could identify not all of genes but only limited genes as being common between two experimental conditions: without and with stress.

**Table 10.** Confusion matrix of coincidence between selected 30 singular value vectors selected among all 1096 singular value vectors, *u*_*ℓj*_, attributed to samples without stress and 24 singular value vectors selected among all 1096 singular value vectors, *v*_*ℓk*_, attributed to samples with stress. For samples without stress, only top 1096 singular value vectors among all 1785 singular value vectors are considered, since total number of singular value vectors attributed to samples without stress is 1096. selected: corrected *P* -values, computed with regression analysis eqs. (12) and (13), are less than 0.01, not selected: otherwise. Odds ratio is as many as 2483 and *P* -values computed by Fisher’s exact test is 1.92× 10^−40^.

|  |  | with stress |  |
| --- | --- | --- | --- |
|  |  | not selected | selected |
| without stress | not selected | 1065 | 1 |
|  | selected | 7 | 23 |

Figure 3 shows the coincidence of selected singular value vectors between samples without and with stress. Singular value vectors with smaller *ℓ*s, i.e., with more contributions, are more likely selected and coincident between samples without and with stress. This can be the side evidence that guarantees that TD based unsupervised FE successfully integrated scRNA-seq data taken from samples without and with stress with avoiding to regard that all are coincident between two samples.

**Figure 3.**
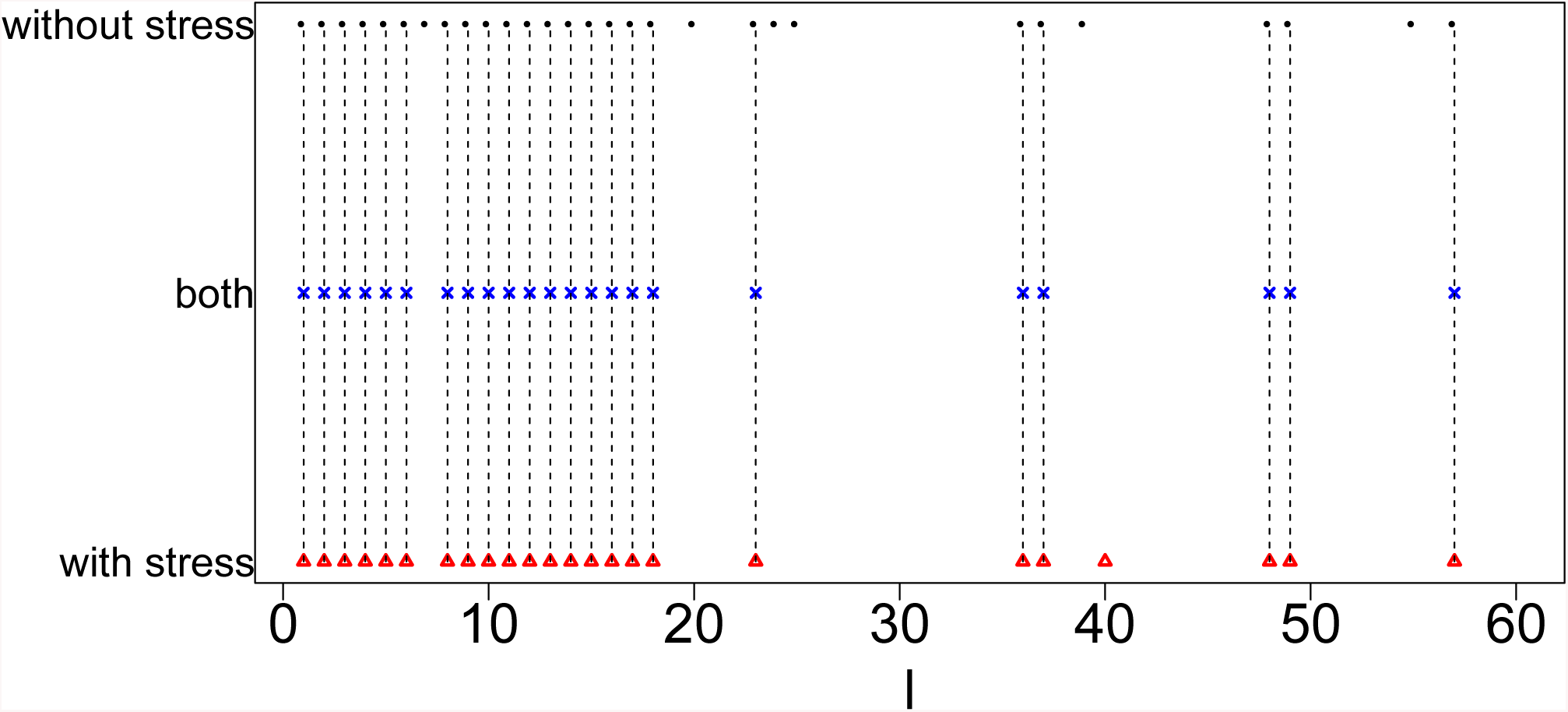
Coincidence between singular value vectors shown in Table 10. Horizontal axis: singular value vector numbering *ℓ*. Black open circles: *ℓ*s selected for samples without stress, blue crosses: *ℓ*s selected for both samples without and with stress, red open triangles: *ℓ*s selected for samples with stress. Vertical black broken lines connect *ℓ*s selected for both samples without and with stress.

Next, we selected genes with following the procedures described in Methods and Materials. The first validation of selected genes is the coincidence between human and mouse. Table 11 shows the confusion matrix that describes the coincidence of selected genes between samples without and with stress. Odds ratio is as large as 270 and *P* -value is 0 (i.e., less than numerical accuracy). Thus as expected, TD based unsupervised FE could not all but limited number of genes associated with cell type dependence.

**Table 11.** Confusion matrix of coincidence between selected 4150 genes for samples without stress and selected 3621 genes for samples with stress among all 24341 genes. Selected: corrected *P* -values, computed with *χ*^2^ distribution that correspond to eqs. (8) and (9) in human and mouse midbrain study, are less than 0.01, not selected: otherwise. Odds ratio is as many as 270 and *P* -values computed by Fisher’s exact test is 0 (i.e. less than numerical accuracy).

|  |  | with stress |  |
| --- | --- | --- | --- |
|  |  | not selected | selected |
| without stress | not selected | 19894 | 297 |
|  | selected | 826 | 3324 |

**Table 12.** Five top ranked terms from ‘GTEx Tissue Sample Gene Expression Profiles up’ by Enrichr for 3324 genes selected commonly between samples without and with stress.

| Term | Overlap | *P*-value | Adjusted *P*-value |
| --- | --- | --- | --- |
| GTEx-WWYW-0011-R10A-SM-3NB35_brain_female_50-59_years | 1006/2885 | $2.7880 \times 10^{-151}$ | $8.135 \times 10^{-148}$ |
| GTEx-T6MN-0011-R1A-SM-32QOY_brain_male_50-59_years | 859/2317 | $2.9865 \times 10^{-144}$ | $4.3575 \times 10^{-141}$ |
| GTEx-QVUS-0011-R3A-SM-3GAFD_brain_female_60-69_years | 963/2759 | $6.8195 \times 10^{-144}$ | $6.6325 \times 10^{-141}$ |
| GTEx-T2IS-0011-R3A-SM-32QPB_brain_female_20-29_years | 967/2792 | $5.5265 \times 10^{-142}$ | $4.0315 \times 10^{-139}$ |
| GTEx-WZTO-0011-R3B-SM-3NMC6_brain_male_40-49_years | 991/2972 | $2.6805 \times 10^{-133}$ | $1.5645 \times 10^{-130}$ |

Finally, we tried to evaulate if genes selected are tissue type specific, i.e., hypothalamus. We have uploaded commonly selected 3324 genes to Enrichr. ‘GTEx Tissue Sample Gene Expression Profiles up’ suggest that all five top ranked terms are brain with high significance (adjusted *P* -vlaues are less than 1× 10^−130^). This suggests that TD based unsupervised FE successfully limited number of genes related to brains even using closely related samples. In order to be more specific, we checked ‘Allen Brain Atlas up’ in Enrichr. Then we found that all top ranked five terms are hypothalamic (Table 13). It is interesting that TD based unsupervised FE could successfully identify hypothalamic specific genes only using scRNA-seq retrieved from hypothalamic. It is usually required to use data taken from other tissues in order to identify tissue specific genes because we need to compare targeted tissues and not targeted tissues in order to identify genes expressive specifically in target tissues. The successful identification of genes specific to something without using the comparison with other samples was also observed previously when tumor specific genes are tried to be identified with TD based unsupervised FE (30). In this sense, TD based unsupervised FE methods are effective not only when genes common between two distinct conditions are sought but also when genes common between two closely related conditions are sought. Thus, it is unlike that the success of TD based unsupervised applied to scRNA-seq is accideintal.

**Table 13.** Five top ranked terms from ‘Allen Brain Atlas up’ by Enrichr for 3324 genes selceted commonly between samples without and with stress.

| Term | Overlap | *P*-value | Adjusted *P*-value |
| --- | --- | --- | --- |
| Paraventricular hypothalamic nucleus | 120/301 | $3.38 \times 10^{-22}$ | $7.41 \times 10^{-19}$ |
| Paraventricular hypothalamic nucleus, parvicellular division | 119/301 | $1.15 \times 10^{-21}$ | $1.27 \times 10^{-18}$ |
| Paraventricular hypothalamic nucleus, parvicellular division, medial parvicellular part, dorsal zone | 117/301 | $1.29 \times 10^{-20}$ | $9.42 \times 10^{-18}$ |
| paraventricular nucleus, cap part | 116/301 | $4.22 \times 10^{-20}$ | $2.31 \times 10^{-17}$ |
| Paraventricular hypothalamic nucleus, magnocellular division | 115/301 | $1.36 \times 10^{-19}$ | $5.96 \times 10^{-17}$ |

## 4 DISCUSSIONS AND FUTURE WORK

In this study, we applied TD based unsupervised FE to the integration of scRNA-seq data sets taken from two species: human and mouse. In the sense of identification of biologically more relevant set of genes, TD based unsupervised FE can outperform PCA based unsupervised FE that previously (34) could outperform three more popular methods: highly variable genes, bimodal genes, and dpFeature. Thus, it is expected that TD based unsupervised FE can do so, too.

For the purpose of integration of two scRNA-seq data set, TD based unsupervised FE has many advantages than other four methods, i.e., PCA based unsupervised FE, highly variable genes, bimodal genes and dpFeature. At first, TD based unsupervised FE can integrate two scRNA-seq data set, not after the selection of genes, but before the selection of genes. This enables us to identify more coincident genes set between two scRNA-seq, in this study human and mouse. As a result, we could identify more coincident results between human and mouse.

The criterion of genes selection is quite robust; they should be dependent upon time points when they are measured. We do not have to specify how they are actually correlated with time. It is another advantage of TD based unsupervised FE.

With applying enrichment analysis to genes selected, we can find many valuable insights about biological process. As a result, we can identify ten key TFs that might regulate embryonic midbrain developments. All of ten selected TFs turn out to be related to brains.

TD based unsupervised FE turn out to be quite effective to integrate two scRNA-seq data set. This method should be applied to various scRNA-seq data sets considering broader scope of investigations.

In future work, we plan to (1) utilize the proposed TD based unsupervised FE under the transfer learning setting; (2) extend the proposed approach to handle the data integration from multiple related tasks; and (3) investigate the performance of the proposed approach when coupled with machine and deep learning algorithms.

## Supporting information

Enrichr for mouse

Enrichr for human

Genes selected for mouse

Genes selected for human

## CONFLICT OF INTEREST STATEMENT

No conflict of interest is declared.

## AUTHOR CONTRIBUTIONS

YHT planed the research, performed analyses, and wrote a paper. TT discussed the results and wrote a paper.

## FUNDING

It was supported by KAKENHI, 17K00417 and 19H05270 and Okawa foundation, grant number 17-10. This project was also funded by the Deanship of Scientific Research (DSR) at King Abdulaziz University, Jeddah, under grant no. KEP-8-611-38. The authors, therefore, acknowledge with thanks DSR for technical and financial support.

## ACKNOWLEDGMENTS

Nothing acknowldegded.

## DATA AVAILABILITY STATEMENT

The datasets analyzed for this study can be found in the GEO.

https://www.ncbi.nlm.nih.gov/geo/query/acc.cgi?acc=GSE76381.

https://www.ncbi.nlm.nih.gov/geo/query/acc.cgi?acc=GSE74672

## Notes

#### Summary of Updates

It is a comprehensive revision for addressing reviewer's comments.

